# Developmental coordination disorder affects the pre-ordering of sequential movements

**DOI:** 10.64898/2026.06.12.731668

**Authors:** Helena Wright-Wieckowski, Kate Wilmut, Katja Kornysheva

## Abstract

Research suggests that motor difficulties in Developmental Coordination Disorder (DCD) are related to altered motor sequence planning, but it is unclear which mechanisms are affected, particularly in adults. This study addresses that gap by examining how the order of upcoming movements during the planning of skilled typing sequences affects motor production in adults with DCD. Previous monkey neurophysiology and behavioural findings in humans have shown that elements of a sequence are pre-ordered prior to execution, known as competitive queuing (CQ). CQ quality is predictive of subsequent performance, with skilled performers, those with fewer errors, having a larger position-dependent difference.

DCD (N=28) and control participants (N=54) performed two 4-element finger sequences from memory in a delayed sequence production task over 3 sessions. Probe trials, which involved participants performing a single press after the Go Cue, assessed motor planning at each sequence position by measuring reaction time (RT) and error rate. We found that adults with DCD had a higher error rate and were slower to initiate and perform correct sequences. In terms of planning, the DCD group showed reduced preordering of sequence elements. Whilst the DCD group had a higher error rate on a working memory task, this was not correlated with the degree of pre-ordering of presses of the upcoming sequence.

These findings suggest that disrupted motor sequence planning in DCD is characterised specifically by a failure to pre-order movements during the retrieval of sequences from memory. Additionally, motor sequence pre-ordering deficits in DCD are independent of general working memory impairments. These results extend prior evidence from motor imagery paradigms, demonstrating that internal modelling deficits are evident during the execution of motor plans.

**Highlights:**

- Adults with DCD show diminished pre-ordering of sequential movements during planning.
- The DCD group were slower to initiate and perform correct sequences from memory.
- Motor sequence planning is distinct from working memory performance.
- This provides evidence for the IMD hypothesis in a skilled sequential task.

## Introduction

Approximately half of the global population relies on typing for communication and digital tool use, an activity that requires the precise coordination of complex movement sequences. However, individuals with Developmental Coordination Disorder (DCD)/Dyspraxia, often experience difficulties with movement coordination, particularly when executing sequenced actions, including typing. These challenges are believed to result from impairments in movement planning and organisation (Bhoyroo et al., 2018). DCD is a prevalent neurodevelopmental disorder affecting around 6% of the population (Blank et al., 2019) and is characterised by difficulties in acquiring fine and/or gross motor skills (Kirby and Sugden, 2007). These motor impairments, which are not attributable to intellectual disability, visual impairment, or neurological conditions, hinder the planning, acquisition, and execution of age-appropriate motor tasks (American Psychiatric Association, 2013). DCD typically emerges in early childhood and often persists into adulthood, with approximately 75% of individuals continuing to experience motor coordination problems such as clumsiness, slowness and inaccuracy of motor tasks that interfere with daily activities such as typing (Barnett & Stuart, 2024; Blank et al., 2019).

Although numerous studies have examined the movement production in children with DCD (Jelsma et al., 2021; Verbecque et al., 2021; Wilson et al., 2013, 2020), and some in adults (Engel-Yeger, 2020; McIntyre et al., 2017; Suzuki et al., 2020). Research investigating the mechanisms underlying motor planning difficulties remains limited. Nevertheless, existing evidence suggests that impairments in motor planning may contribute to the deficits observed in DCD (Bhoyroo et al., 2018, 2019) but few studies have examined these mechanisms in adults. End-state comfort (ESC) tasks involve the ability to plan a movement sequence such that the final step results in a comfortable posture, which may involve adopting a starting posture that is uncomfortable (Wilmut & Byrne, 2014). Studies on grip selection in children with DCD using ESC tasks revealed that as task complexity increased, children with DCD often showed poorer performance compared to their peers (Bhoyroo et al., 2020). Children with DCD chose fewer grips for ESC and instead favoured initial grips with minimal rotation compared to their peers (Wilmut & Byrne, 2014), indicating impaired anticipatory motor planning, as they prioritised immediate comfort over future task efficiency. A longitudinal study that used an ESC task to assess motor planning differences in DCD, compared to controls, and found developmental delay (Adams et al., 2017). At baseline, children with DCD were slower and less accurate than their peers, whilst both groups showed similar rates of improvement in action planning over two years, the DCD group remained slower and made significantly more errors (Adams et al., 2017). In adults with DCD, grip selections were often used if they required a minimal initial rotation grasp during ESC tasks that required more complex movement sequences (Wilmut & Byrne, 2014). This preference suggests ongoing difficulties with planning complex motor actionsthat persists into adulthood. One prominent theoretical account that may explain the deficit in motor planning is the internal modelling deficit hypothesis (IMD). The IMD proposes that individuals with DCD have difficulty generating accurate internal models. An internal model is a motor representation that is used to predict and control movement, which provides stability to the motor system by predicting the outcome of movements before sensory feedback is available (Miall & Wolpert, 1996; Wolpert et al., 1995). An inaccurate forward model, may contribute to the difficulties in learning new action sequences (Adams et al., 2014; Wilson, Smits-Engelsman, et al., 2017).

Hand rotation tasks (HRT) have been used to examine the internal representation of movement, as participants have to imagine the rotation of the hand (Hyde, 2014; Hyde et al., 2018). When stimuli are presented at varying orientations, performance tends to decline, becoming slower and less accurate as the required hand rotations become more complex (Barhoun et al., 2021). When performing a HRT, DCD participants were found to be significantly less efficient than controls, but there were no performance differences between groups on the letter number rotation task (Barhoun et al., 2021). This task only requires visual imagery, therefore suggesting that the deficit is linked to motor imagery specific tasks (Barhoun et al., 2021). Furthermore, previous studies have found that adults with DCD engaged in motor imagery (MI) to complete a hand rotation task/hand laterality task, finding they were significantly less efficient at the task (Hyde, 2014; Hyde et al., 2018). Additionally, when performing the HRT task in the scanner, adults with DCD showed lower %BOLD signal change with increasing angles of rotation compared to controls in the occipito-parietal and parieto-frontal networks, as well as in the cerebellum. Behaviourally, the probable DCD (pDCD) group demonstrated lower accuracy and more variable performance. (Kashuk et al., 2017). These brain regions are primarily involved in motor planning, sequence representation and the execution of automatic skilled movements (Culham & Valyear, 2006). Dysfunction in these areas could lead to the use of compensatory strategies or a dependence on slower feedback-based control (Wilson, Smits-Engelsman, et al., 2017).

Various studies have linked differences in brain areas such as the cerebellum, parietal and prefrontal cortices with the deficits in DCD (Kashiwagi et al., 2009; Kashuk et al., 2017; Wilson, Smits-Engelsman, et al., 2017; Zwicker et al., 2011), areas primarily involved in movement planning (Culham & Valyear, 2006; Freedman & Ibos, 2018; Spencer et al., 2007; Wilson, Smits-Engelsman, et al., 2017). Furthermore, children with DCD showed reduced activation in the posterior parietal cortex (PPC) and postcentral gyrus, regions involved in integrating multimodal motor information (Freedman & Ibos, 2018), sequencing skilled movements (Yewbrey et al., 2023), and generating internal motor representations during a visuomotor tracking task, compared to controls (Kashiwagi et al., 2009). Therefore, these findings support the view that motor planning of complex movements is disrupted in DCD, potentially due to impaired internal motor representation, particularly for sequences of actions.

Therefore, these findings suggest that adults with DCD may have a weaker internal action representation when performing sequences of movement, consequently they may rely more on feedback to complete the movement (Adams et al., 2014). So far, much of the existing research has focused on motor imagery tasks to investigate internal movement representation (Adams et al., 2017; Barhoun et al., 2021; Hyde, 2014; Hyde et al., 2018; Kashuk et al., 2017). Motor imagery primarily taps into the ability to mentally stimulate individual actions without performing these movements, whereas the planning of skilled movement sequences involves a different and potentially more complex aspect of motor planning, one that integrates sequencing, timing and the coordination of multiple actions. This distinction is important, as complex movements are often impaired in individuals with DCD (Fuelscher et al., 2016; Gentle et al., 2021; Wilmut & Byrne, 2014). The present research therefore aims to examine motor planning in the context of skilled action sequences, to better understand how planning deficits contribute to the broader difficulties with movement execution in DCD.

Animal and human neurophysiology studies of movement sequence planning have demonstrated that movements are planned competitively in parallel prior to execution, queued in order of the upcoming sequence (Averbeck et al., 2001; Kornysheva et al., 2019; Mantziara et al., 2021). This also supports computational models of competitive queuing (CQ) (Burgess & Hitch, 2005; Houghton, 1990; Houghton & Hartley, 1996). These findings have since been replicated behaviourally using probe trials (Mantziara et al., 2021). In these trials, participants were trained to perform sequences from memory in a delayed sequence production paradigm. During probe trials, they were shown a sequence cue that triggered sequence planning but instead they were asked to make only a single, guided keypress corresponding to a specific position within the sequence. Responses to earlier sequence positions typically resulted in faster reaction times and fewer errors, reflecting a positional gradient that mirrors the parallel planning processes observed in neural activity (Mantziara et al., 2021). Furthermore, the strength of the CQ gradient has been linked with more accurate performance (Kornysheva et al., 2019), and faster RTs (Mantziara et al., 2021), showing it to be a marker of skilled sequence performance.

We aim to explore whether difficulties in motor planning are linked to limitations in working memory, or whether they reflect a distinct impairment in skilled motor memory. Previous research has found that children with DCD were less proficient in working memory tasks compared to controls, particularly in the visuospatial domain (Alloway & Temple, 2007). Additionally, we introduced an inhibition task (Go/No-Go), as previous research has shown that young adults with DCD have difficulty engaging inhibitory mechanisms necessary for effective motor control (He et al., 2018). Given that motor planning involves not only the preparation of intended actions but also the suppression of competing or premature responses, deficits in inhibition may interfere with the effective organisation of action sequences.

Here, we investigated sequential motor planning to identify whether the primary deficit in motor planning in DCD lies in the initial pre-ordering of ordinal sequence structures prior to movement execution, and whether this underlies the prolonged initiation times and heightened error rates observed in this group.

### Methods and materials Participants

Participants were recruited via the research participation scheme (RPS), the Centre for Human Brain Health (CHBH) mailing list, Oxford Brookes DCD mailing list, Dyspraxia Foundation (DF) and online via social media. All participants were right-handed, aged between 18 and 35, with normal hearing and normal or corrected-to-normal vision, who were proficient in the English Language and had no finger, hand or wrist injuries and no prior history of neurological disorders. Participants received financial reimbursement at a rate of £3.50 per session, or £10.50 for all three sessions. For non-financial reimbursement, participants received 1 SONA credit/hour. All participants gave informed consent according to the Ethics Committee at the University of Birmingham.

A total of 182 participants initially signed up for the study: 140 controls and 37 DCD participants. When participants registered for the study, they indicated which group they belonged to, i.e. Volunteer group A: Individuals with clinically diagnosed dyspraxia/DCD, but no prior history of neurological disorders and finger, hand or wrist injury. Volunteer group B: Individuals who suspect they have dyspraxia/DCD but no prior history of neurological disorders and finger, hand or wrist injury. Volunteer group C: Volunteers without prior history of dyspraxia/DCD, psychiatric or neurological disorders and finger, hand or wrist injury. This process initially put individuals into groups and then we defined each group as either belonging to the control or DCD group by using the following methods. There were too few participants in volunteer group B and as such this group was not used in further analyses.

The ADC checklist (Kirby et al., 2010), is the first screening tool developed for adults to identify difficulties experienced by those with DCD. This is divided into 4 subscales; subscale A was designed to make sure symptoms were present in childhood, whilst subscale B, C and D examine current functioning. Following the scoring guidelines in Kirby et al (2010), a combined score on subscale A of >17 and an overall score of 56 indicate probable DCD; therefore, we excluded any control participants that scored above this threshold. We used the adult dyspraxia checklist (ADC) to inform our participant exclusion criteria for two reasons: firstly, we wanted to exclude control participants that had probable DCD and secondly, we used the ADC checklist as an additional measure for our adult participants to further assess whether they met the DSM-5 criteria.

Furthermore, for inclusion in the DCD group, it required that participants self-reported that they had a formal clinical diagnosis of DCD and they were further screened according to the DSM-5 criteria for DCD, which requires: (A) significantly impaired acquisition and execution of coordinated motor skills relative to age expectations, (B) persistent motor difficulties that interfere with daily life and academic functioning, (C) symptom onset in early development, and (D) motor deficits not better explained by intellectual disability, visual impairment, or a neurological condition (Kirby et al., 2010). To address part D, in the questionnaire participants were asked if they had a diagnosis of any of the following disorders: a) DCD, Developmental Dyslexia (DD), Specific language impairment (SLI), Attention deficit hyperactivity disorder (ADHD), Autism spectrum disorder (ASD), Asperger’s syndrome, Developmental stuttering/stammering, Tic/Tourette’s, Intellectual development disorder (IDD), b) Visual impairment (not corrected by visual aids), Cerebral Palsy, Muscular dystrophy, Neurodegenerative disorder e.g. (Alzheimer’s disease, Ataxia, Huntington’s disease, Parkinson’s disease, Motor neuron disease, Multiple system atrophy, Progressive supranuclear palsy), Traumatic brain injury (TBI), Stroke or other. If either control or DCD participants reported that they had any neurological disorders (b), they were excluded from the study and control participants were excluded if they had any of the above disorders (a or b). Additionally, control participants were excluded if they scored high (>56) on the ADC checklist, indicating they could have DCD (Figure 1). This was because any of these additional disorders or scoring highly on the ADC could influence movement and affect the results.

**Figure 1.**
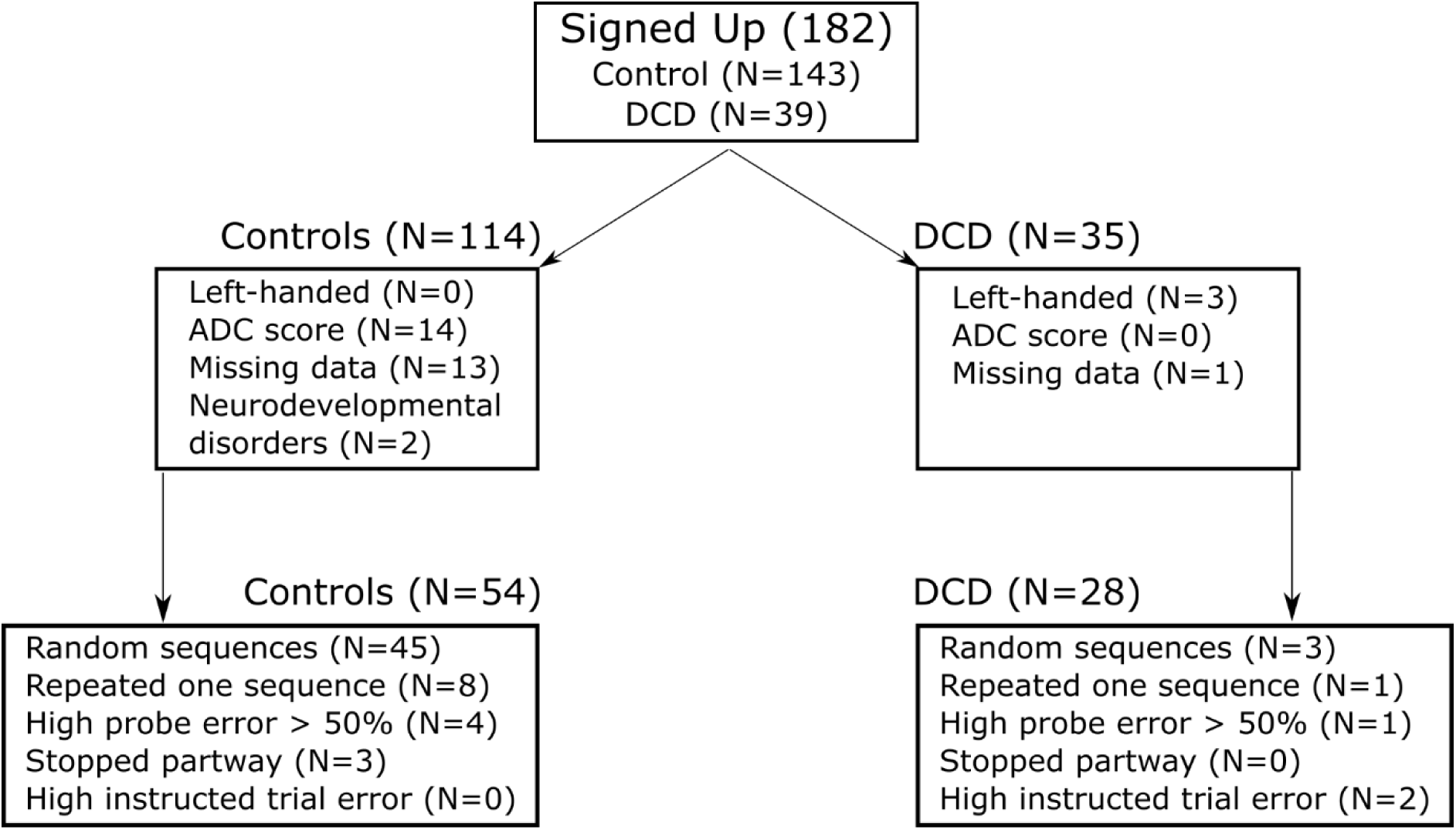
Diagram to show the process of excluding participants.

Participants were then excluded from both the Control and DCD groups if they were left-handed or had missing data (i.e., one or more data collection sessions were not completed). Furthermore, participants from both groups were excluded if they performed random sequences, either because they made random key presses or because they failed to associate the two motor sequences with their respective visual cues. Participants were also excluded if they completed only one of the sequences, stopped partway through the task, had a high error rate on instructed trials, or had a high probe error rate (>50%), resulting in an insufficient number of correct trials for analysis.

Finally, we had 82 participants, of whom 54 were control participants and 28 self-reported a diagnosis of DCD. 22 participants had DCD, 3 had DCD and ADHD, 1 had DCD, ADHD and developmental dyslexia (DD), 1 had DCD, DD and Developmental language disorder (DLD), 1 had DCD and Autism spectrum disorder (ASD). All participants were right-handed and the Edinburgh handedness inventory was used to assess this (Oldfield, 1971). There were no differences between the handedness scores for groups, t(43.18) = 1.20, *p* = .24. control (*M* = 83.43, *SD* = 22.90), DCD (*M* = 75.54, *SD* = 30.53). Participants were aged between 18-35 years old. The DCD group (*M* = 26.93, *SD* = 5.41) was older compared to controls (*M* = 22.04, *SD* = 4.03), t(45.080) = -4.27, *p* < .001.

### Apparatus and stimuli

Participants completed the study online using Pavlovia (www.pavlovia.org) on a computer or laptop from the comfort of their home. They were instructed to sit in a quiet place, without distractions and complete the task with the sound on. Participants placed their right index, middle, ring and little fingers on the y,u,I,l buttons, respectively.

The abstract images used were created by applying a sine-wave function on a radial space and were individually created by varying their curvature and frequency. These images were similar to those used in Helie & Cousineau (2005). Each trial began with an abstract image (sequence cue) that had a fixed duration of 500ms, followed by 500ms during which a fixation cross was shown (Figure 2). Subsequently, a black right-hand stimulus appeared as a Go cue. For instructed trials, the background of the Go cue was grey with a digit cue indicating which finger participants needed to press, and in memory trials, the background was green, and there were no digit cues as participants produced the sequence from memory. After participants performed the movement, a fixation cross was shown for 1000ms, and then participants received feedback, which was shown for 1000ms. If the sequence position was correct, the feedback consisted of a cross (x) and for incorrect trials, a dash (-) was shown. In addition to this, participants were shown points ranging from 1 to 10 with an image, depending on how quickly participants produced the movement. Probe trials were introduced to measure movement availability during sequence planning. Probe trials started the same as sequence trials, beginning with a fractal cue (500ms), followed by a fixation cross (500ms). A probe cue was then presented, prompting participants to make a single key press corresponding to a specific position in the sequence. Participants received feedback for these trials of up to 5 points.

**Figure 2.**
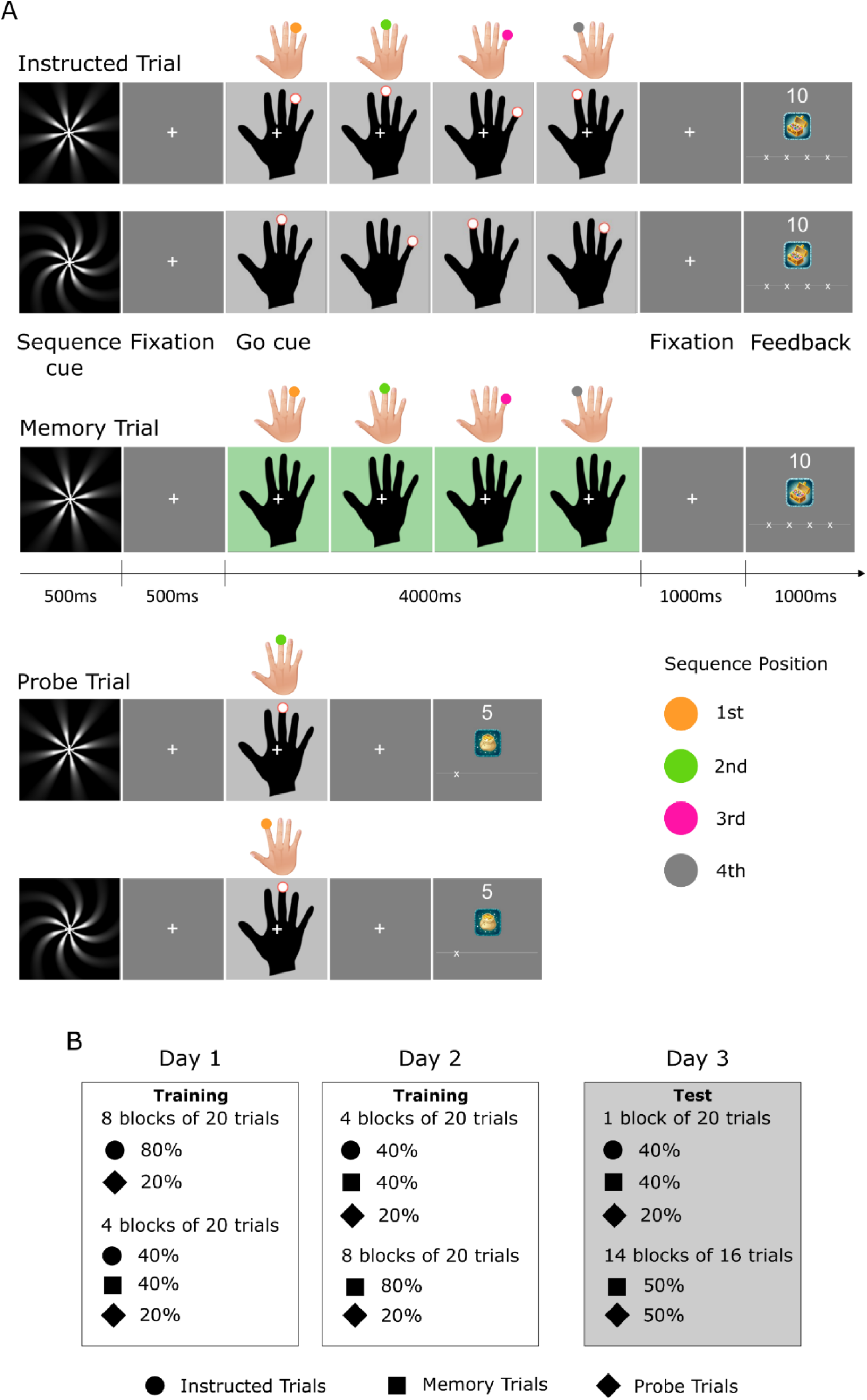
**A.** This shows the diagram of the trials, instructed, memory and probe trials. For probe trials the sequence position indicates which position of the sequence that is being probed. **B.** The trials included for each stage of the experiment: training included day one and two, whilst the test phase included day three.

### Procedure

The experiment took place across 3 days, including 2 training days (day 1 and day 2) and 1 test day which typically took 30 minutes to complete. At the beginning of the study, participants watched an instruction video. Following this, their understanding of the task instructions were tested through three questions. Participants needed to get all 3 questions correct to move on to the task. If they did not get all three questions correct, they could retake the quiz. Day 1 consisted of 8 blocks of 20 trials (80% instructed and 20% probe), 4 blocks of 40% instructed, 40% memory-guided and 20% probe. Day 2 had 4 blocks of 40% instructed, 40% memory-guided and 20% probe and then 8 blocks of 80% memory-guided and 20% probe. Day 3 started with a short refresher block and then 50% probe trials and 50% memory-guided trials. After completing the finger press task, participants completed the ADC checklist, n-back task to assess working memory capacity and the go/no-go task to assess inhibition.

For the n-back task, participants completed three levels of difficulty (1-back, 2-back, 3-back), which required recalling the spatial location of a black square that appeared on a grey square grid, either one, two or three trials earlier, respectively. Similarly, for the go/no-go task, there were two within-subject conditions based on stimulus frequency: the target stimulus (“X”) was presented either 80% or 20% of the time.

### Data analysis

Data analyses were performed using MATLAB (The MathWorks Inc. (2024). *MATLAB version: 9.15.0 (R2024a)*. Natick, Massachusetts: The MathWorks Inc. https://www.mathworks.com), and RStudio (*Posit team (2025). RStudio: Integrated Development Environment for R.Posit Software, PBC, Boston, MA. URL* http://www.posit.co/).

#### Sequence production

All trials for analyses were taken following the refresh block on day three of the study. This part of the task included only memory and probe trials. At this point of the task, the participants are producing the sequences from memory.

Due to a coding error, a subset of participants completed 2 fewer training blocks compared to other participants on day one of the study. This included 18 participants in the control group and 22 participants in the DCD group. The two additional training blocks on the first day of the experiment included 8 instructed trials per sequences and 4 probe trials in each block. To determine the effect of this, we used ‘training’ as a covariate in later analyses, which included two groups. Those that had less training or more training on day 1.

To measure performance differences between groups we investigated sequence initiation time, which was defined from the Go cue to when participants completed the first key press. Sequence movement time (MT), defined as from the first key press to the last key press. Error rate, defined as an incorrect button press and calculated as a percentage. An ANCOVA was used to investigate the effect of group (control vs DCD) on performance measures (initiation time, MT, error rate) while controlling for the covariates training and age (Figure 3).

**Figure 3.**
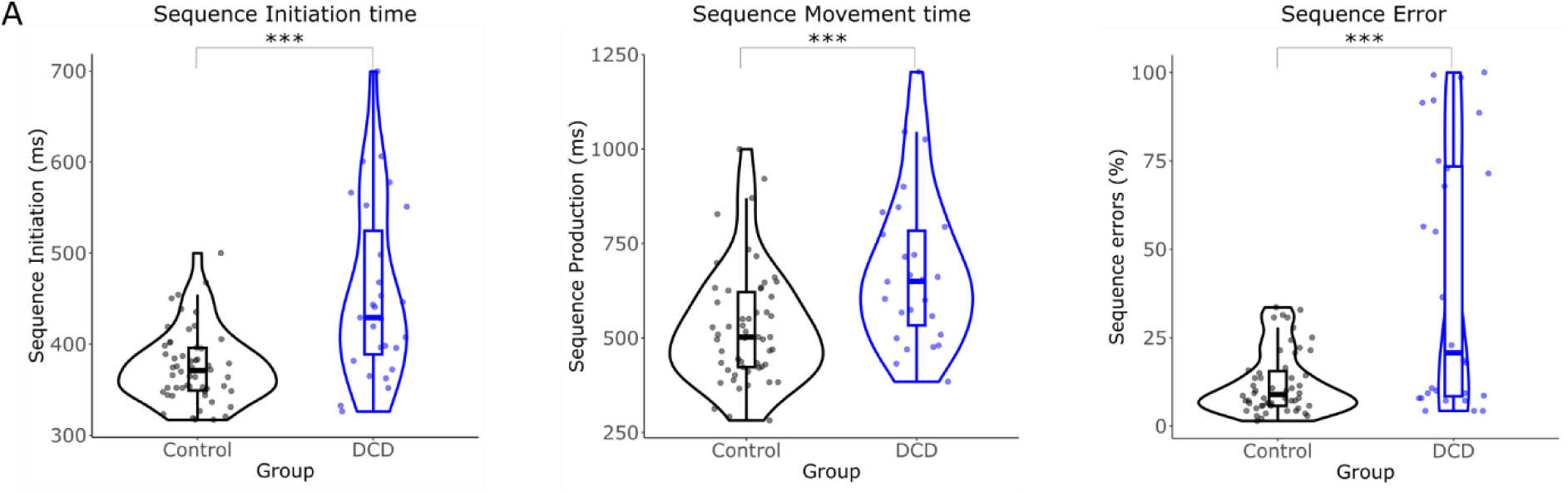
The sequence production results from the test phase for correct trials include the median initiation time (measured from the go cue to the first position press), the movement time (from the first position press to the last), and the percentage of incorrect sequences.

To assess group differences in working memory, we conducted a mixed design ANCOVA to examine the effects of group and nbackType (nback1, nback2, nback3) on performance in the nback task, controlling for age. Pairwise comparisons we conducted for the group x nbackType interaction using holm-adjusted post hoc tests (Figure 7a). Post hoc pairwise comparisons were conducted using estimated marginal means adjusted for age, ensuring that all comparisons accounted for the covariate. An ANCOVA was also performed on the inhibition task with the 20% and 80% conditions (Figure 7b).

#### Sequence planning

To measure movement availability during planning, we used the median RT of correct trials and the error rate of probe trials at each sequence position (1^st^, 2^nd^, 3^rd^, 4^th^) as our dependent measures (Mantziara et al., 2021).

A mixed model ANOVA design was used to investigate whether there were differences between group and position in probe trials using raw RT’s and error rates. Significant interaction effects were further explored using one-way ANOVAs to examine group differences at each position, and paired t-tests to assess position differences within each group. Interaction effects between positions (2-1, 3-2, 4-3) were assessed within each group and then compared between groups to find the difference of differences. Additionally, the percentage increase in RT and error rate relative to the first position for each participant was calculated, so the position-dependent differences could be visualised (Figure 4). Following this, we calculated the relative differences for both RT and error rate between adjacent positions (the second minus the first position, the third minus the second position and the fourth minus the third position). This was done to be able to quantify sequence planning into a single value which was used in further analyses. A large value, indicating an increase from the first position to the second, second to the third and third to fourth suggested that there was a larger position-dependent difference, indicating that movements were pre-planned prior to sequence execution.

**Figure 4.**
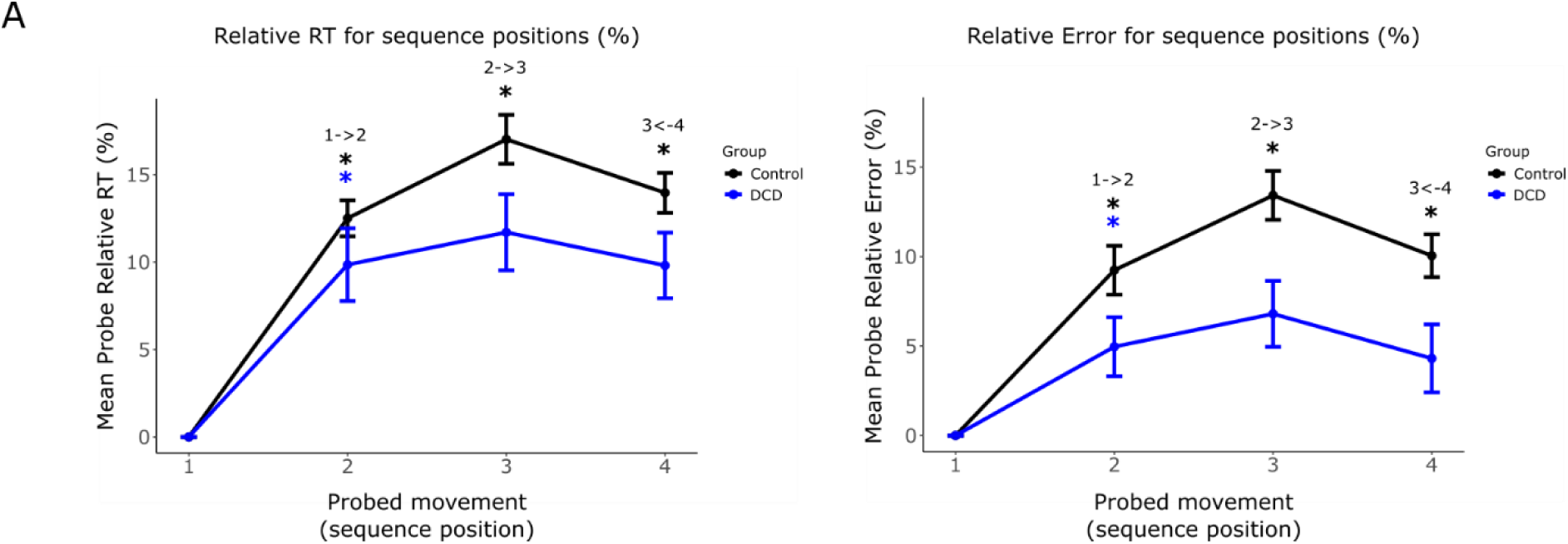
The relative RT and error to the first position for each sequence position for probe trials for the DCD group (blue) and the control group (black). Within-group increases between each position are shown as follows: control (black), DCD (blue).

**Figure 5.**
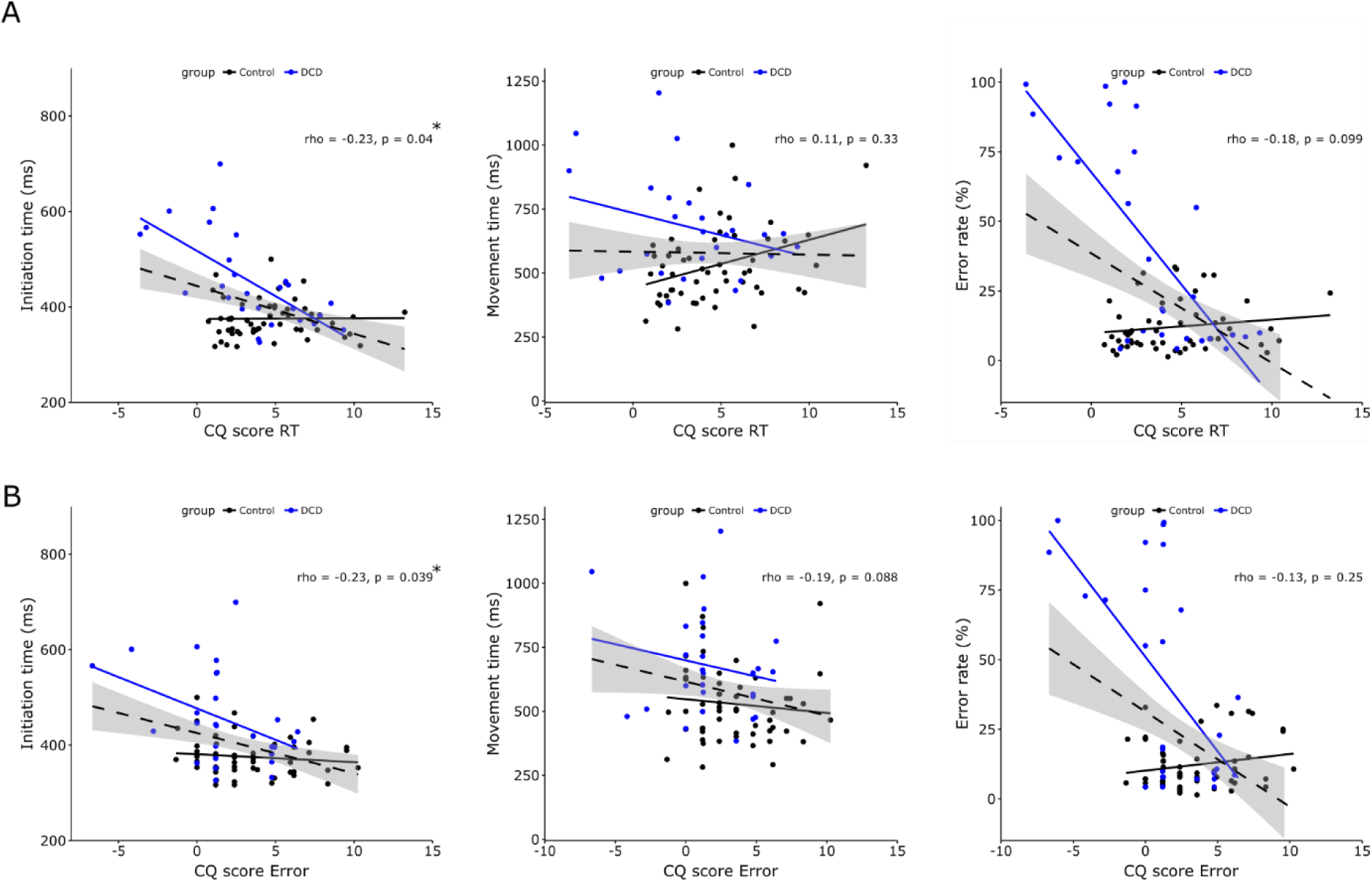
**A.** Scatter plots showing planning (CQscoreRT) and the relationship with the production measures (Initiation time, movement time, error rate) and B. CQscoreError. The DCD group (blue), Control group (black) and the correlation including both groups (dashed).

We subsequently performed a mediation analysis using the PROCESS macro in R (model 4) configured for a binary dependent variable, testing the probability that the relationship between production measures (sequence initiation time, MT and error rate) and a diagnosis of DCD (group)_was mediated by planning measures (CQscoreRT and CQscoreError) (Figure 6). This used a logistic regression for the form of the conditional model and utilised bootstrapping to construct bias-corrected confidence intervals for the indirect effect.

**Figure 6.**
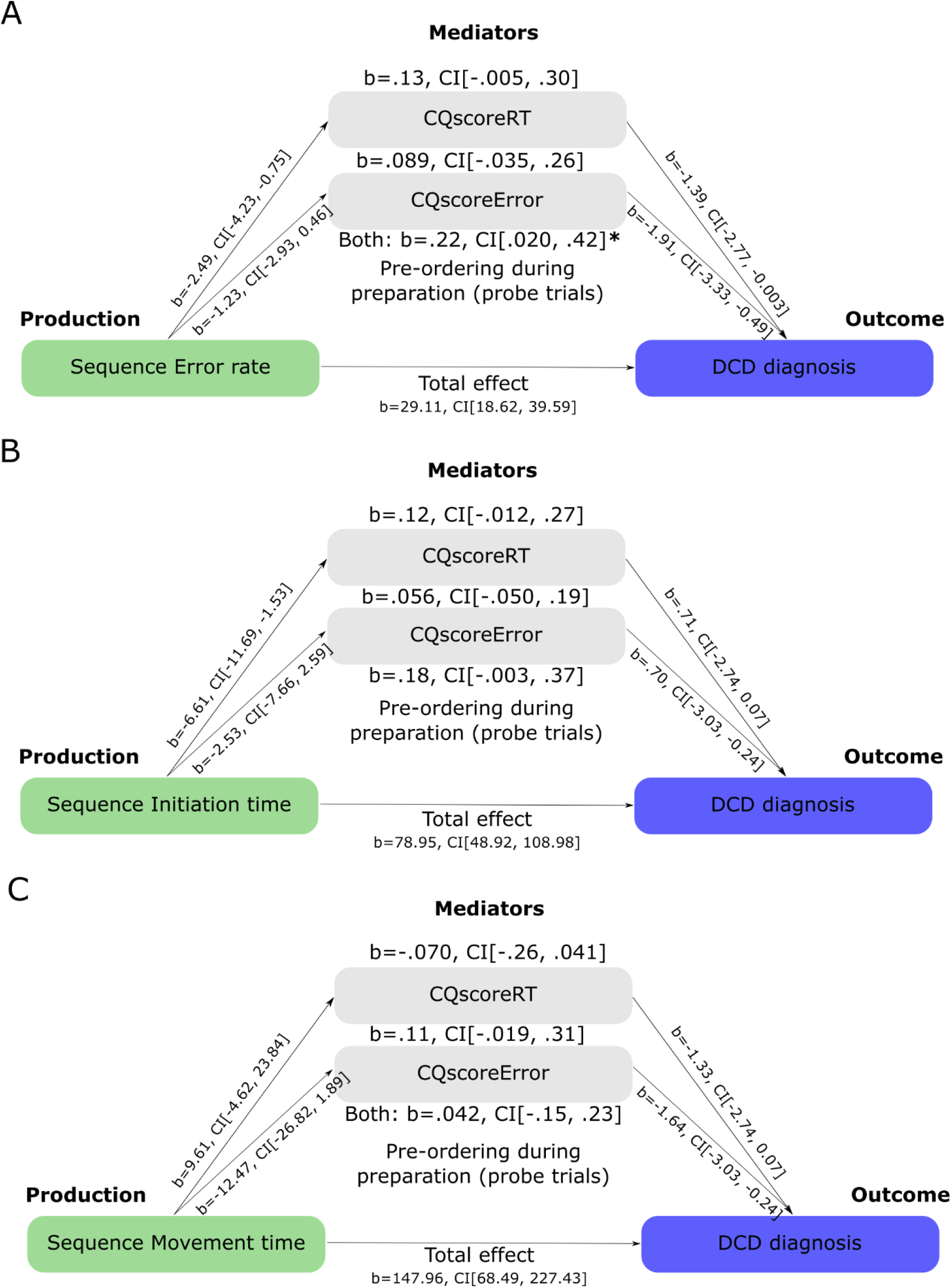
This shows the mediation analysis diagram to examine how production (**A.** Sequence initiation time, **B.** Movement time, **C.** Error rate) predicts DCD diagnosis, with mediators planning (CQscoreRT, CQscoreError).

**Figure 7.**
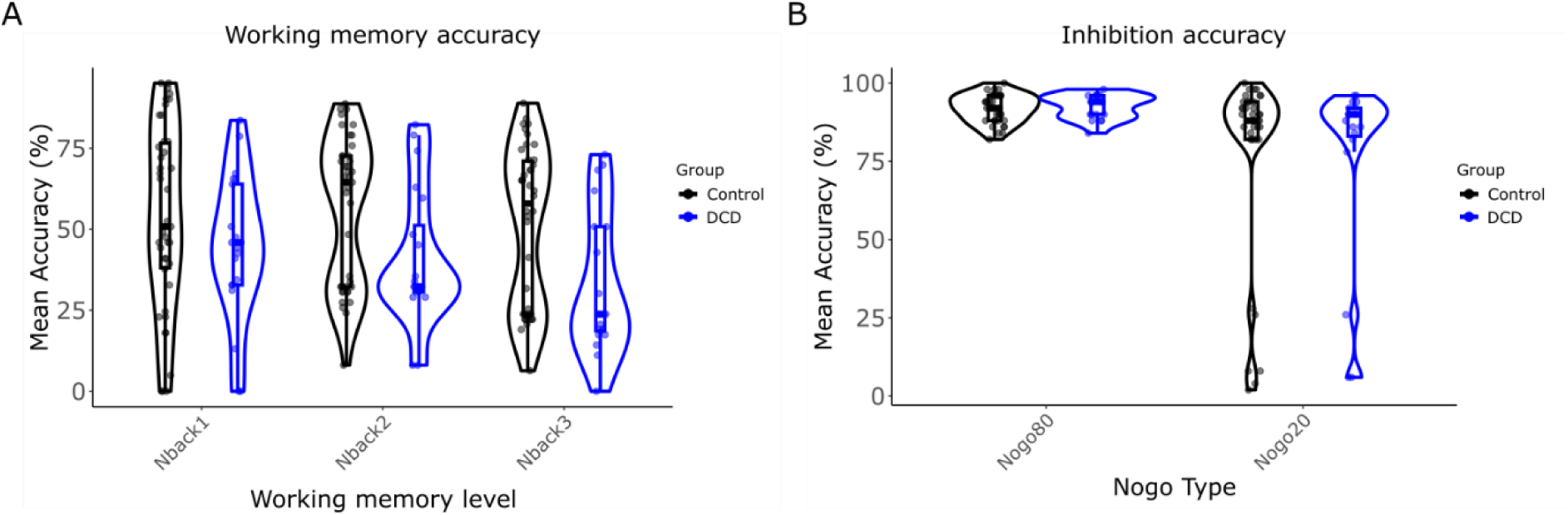
N-back (working memory) task and go-nogo (inhibition) task. **A.** The accuracy percentage for control participants (black) and DCD participants (blue) for each level of the nback task. **B.** The accuracy percentage for each type of the nogo task 80% common and 20% rare for both groups.

**Figure 8.**
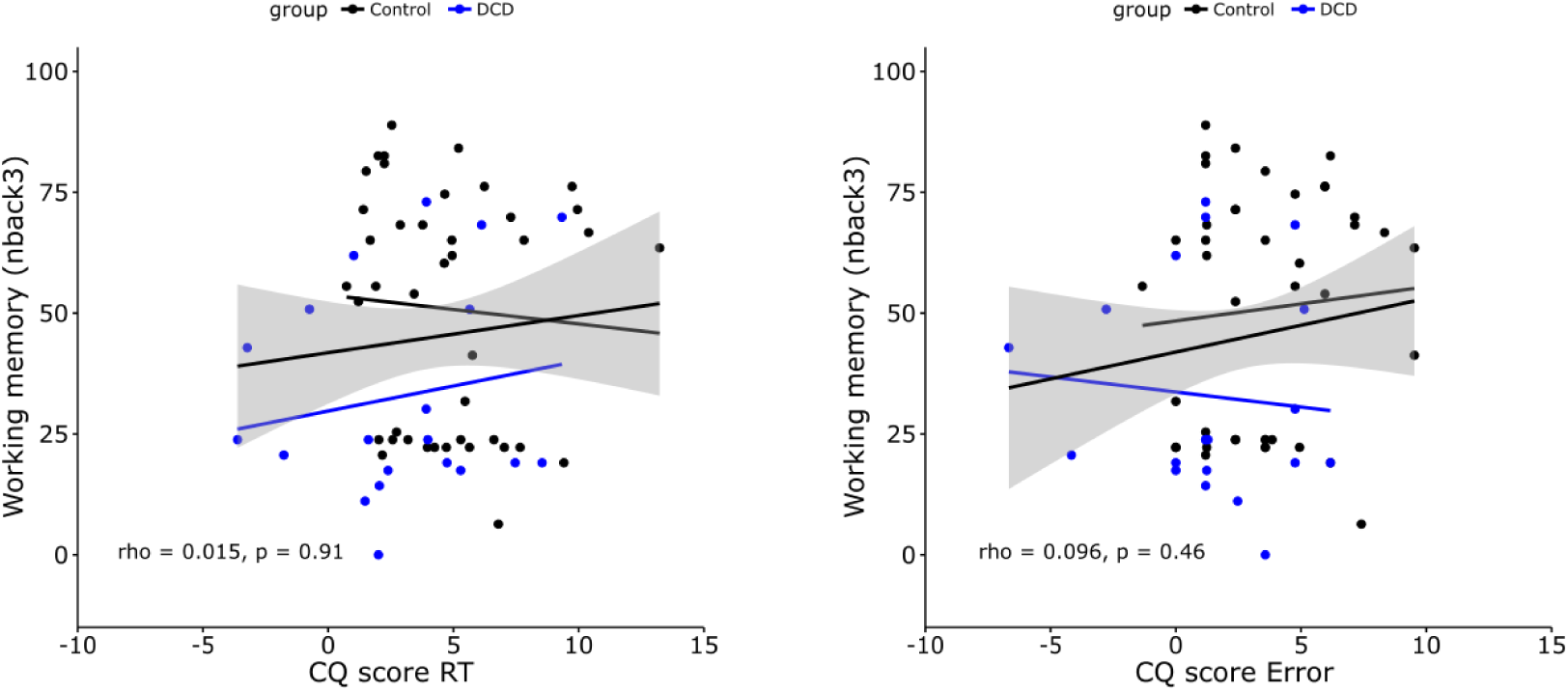
Scatter graphs showing the correlations between planning (CQscoreRT/CQscoreError) and working memory (nback3).

To further investigate the relationships between planning (CQscoreRT/Error), performance measures (sequence initiation time, movement time, error rate), working memory (nback1, nback2, nback3), inhibition (nogo20, nogo80) and ADC score, we conducted a permutation-based correlation analyses with Spearman’s rho with 5000 iterations (Figure 9). This was done to explore how these factors interacted to gain a better understanding of the influences on planning.

**Figure 9.**
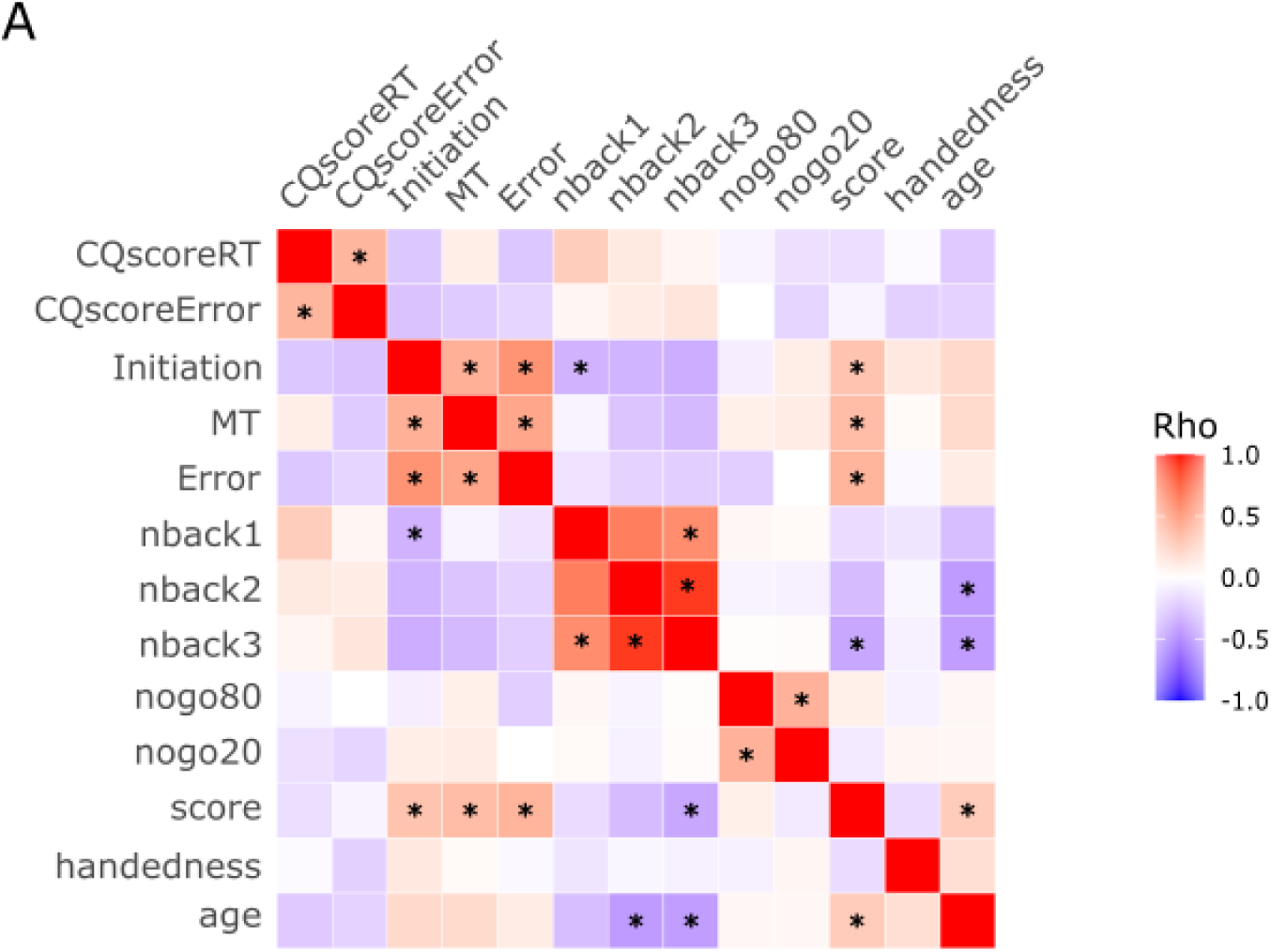
This correlation matrix shows motor planning (CQscoreRT/Error), sequence performance measures (initiation time, movement time, error rate), working memory (nback1,2,3), inhibition (go-nogo 80%, 20%) and ADC overall and subscale A score. Significant Spearman correlations after permutation-based FDR correction (p<.05) are marked with *.

#### Statistical analysis

To assess the assumption of normality for the production data, the residuals from the ANCOVA model were tested using the Shapiro-wilk test. If the residuals were not normally distributed, the dependent variable was transformed using either a logarithmic or square root transformation to better meet the assumptions of ANCOVA. For the probe trial analysis, we tested the assumption of sphericity using Mauchly’s test, when this was violated Greenhouse-Geisser corrections were applied.

## Results

We investigated whether the DCD and control groups differed in performance measures of sequence initiation, movement time, and accuracy. Overall, we found that the DCD group had impaired performance of the sequences from memory compared to controls.

We found that the DCD group were slower to initiate correct sequences (M = 454ms, SD = 95.8) compared to the control group (M = 375ms, SD = 40ms), F(1, 77) = 26.20, p < .001, with no effect of training (F(1,77) = .53, p = .47, and no effect of age (F(1,77) = .18, p = .68. Moreover, the DCD group had a slower MT (M = 678ms, SD = 200ms) compared to controls (M = 530ms, SD = 152ms), F(1,77) = 13.13, p < .001. with no effect of training F(1,77) = .62, p = .43, and no effect of age F(1,77) = .042, p = .84. The DCD group had a higher error rate (M = 41.3%, SD = 36.9%), compared to controls (M = 12.1%, SD = 8.92%), F(1,78) = 25.30, p < .001. There was no effect of training, F(1,78) = 1.12, p = .29, and no effect of age, F(1,78) = 2.29, p = .14. Therefore, this suggests that, as expected, the DCD group tended to have a slower and less accurate performance, and this was not influenced by training or age.

### Individuals with DCD show a reduced competitive queuing of upcoming presses prior to execution

We examined whether the competitive queueing of movements during planning differed between the controls and the DCD group.

Probe RT increased across sequence positions, *F*(2.16, 172.88) = 87.72, *p* < .001, η²ₚ = .52. This effect was not dependent on experimental group, *F*(2.16, 172.88) = 1.98, *p* = .14, η²ₚ = .024. Group differences were observed at Position 1, with the control group (*M* = 450.93 ms, *SD* = 39.95) demonstrating faster responses than the DCD group (*M* = 564.86 ms, *SD* = 281.01); *p* = .016. Similar group differences were found at Position 2, control (*M* = 506.41 ms, *SD* = 47.25) and DCD (*M* = 609.21 ms, *SD* = 260.94); *p* = .018; Position 3, control (*M* = 526.61 ms, *SD* = 56.24) and DCD (*M* = 619.47 ms, *SD* = 266.65), *p* = .030; and Position 4, control (*M* = 513.28 ms, *SD* = 53.64) and DCD (*M* = 613.36 ms, *SD* = 288.92); *p* = .03. Across sequence positions, both groups showed an increase in probeRT from Position 1 to Position 2. The control group increased from *M* = 450.93 ms (*SD* = 39.95) to *M* = 506.41 ms (*SD* = 47.25); *p* < .001, and the DCD group increased from (*M* = 564.86 ms, *SD* = 281.01) to (*M* = 609.21 ms, *SD* = 260.94); *p* < .001. From Position 2 to Position 3, only the control group showed a significant increase, from (*M* = 506.41 ms, *SD* = 47.25) to (*M* = 526.61 ms, *SD* = 56.24); *p* < .001, whereas the increase in the DCD group from (*M* = 609.21 ms, *SD* = 260.94) to (*M* = 619.47 ms, *SD* = 266.65) was not significant, *p* = .17. Similarly, from Position 3 to Position 4, the control group showed a significant decrease, from (*M* = 526.61 ms, *SD* = 56.24) to (*M* = 513.28 ms, *SD* = 53.64); *p* < .001, whereas the DCD group showed no change, decreasing from (*M* = 619.47 ms, *SD* = 266.65) to (*M* = 613.36 ms, *SD* = 288.92); *p* = .82. The absolute probe RT values were also modulated by group, *F*(1, 80) = 7.29, *p* = .008, η²ₚ = .083.

When examining the contrasts between groups, we found no significant difference between the difference of position 2 minus position 1 between groups, p = .21. Or for position 3 minus position 2, p = .26, or for position 4 minus position 3, p = .41.

We found that the probeError increased across sequence positions, F(2.72, 217.76) = 39.07, *p* < .001, η²ₚ = .33, which was dependent on group, F(2.72, 217.76) = 4.14, *p* = .009, η²ₚ = .049. There was only a difference between groups for position 1 (control, *M* = 4.73, *SD* = 2.25), DCD (*M* = 8.37, *SD* = 6.86); *p* = .002. There was an increase from position 1 to 2 for the control group (*M* = 4.73, *SD* = 2.25) to (*M* = 13.54, *SD* = 9.61); *p* <. 001, and also for the DCD group (*M* = 8.37, *SD* = 6.86) to (*M* = 13.30, *SD* = 5.84); *p* = .005), this was the same from position 2 to 3 for the control group (*M* = 13.54, *SD* = 9.61) to (*M* = 17.56, *SD* = 9.34); *p* <. 001) but there was no difference for the DCD group (*M* = 13.30, *SD* = 5.84) to (*M* = 15.02, *SD* = 6.97); *p* = .37. There was a decrease from position 3 to 4 for the control group (*M* = 17.56, *SD* = 9.34) to (*M* = 14.30, *SD* = 8.48); *p* = .010, but not for the DCD group (*M* = 15.02, *SD* = 6.97) to (*M* = 12.81, *SD* = 6.22); *p* = .20. The absolute probeError values were not modulated by group, F(1,80) = .014, *p* = .97, η²ₚ = .00017. Taken together, these results suggest that the DCD group had a smaller position-dependent difference between positions compared to the control group.

When examining the contrasts, we found that the difference between position 2 and position 1, when compared between groups, was significant, p = .04. However, this wasn’t significant when comparing position 3 to position 2, p = .22, or when comparing groups for position 4 minus position 3, p = .58.

In sum, we found that the CQ gradient was less pronounced in the DCD group, suggesting that adults with DCD have a motor planning deficit, as upcoming movements are less pre-ordered compared to controls.

Next, we examined where the performance measures (initiation time, movement time, error rate) correlated with motor planning, specifically the degree of pre-ordering of the upcoming presses (CQscoreRT/Error) (Figure 5). Previously, Mantziara et al (2021) found that performance measures were linked to CQ scores. Replicating these findings, participants who initiated correct sequences more quickly exhibited stronger CQ effects (CQscoreRT: rho = −.25, p = .021; CQscoreError: rho = −.26, p = .02). Other associations were less consistent, consistent with previous work. Participants with shorter movement times tended to show more pronounced CQ, although this did not reach significance and only trended towards significance for CQscoreError (rho = −.22, p = .051), with no relationship observed for CQscoreRT (rho = .08, p = .49). Furthermore, a higher error rate was associated with weaker CQ, which approached significance for CQscoreRT (rho = −.21, p = .059), but not for CQscoreError (rho = −.16, p = .16). Furthermore, we wanted to examine the relationship between motor planning and group. For this we conducted a bivariate correlation for CQscoreRT, t(81)=-2.14, p=.035, cor=-.23 and CQscoreError, t(81)=-2.89, p=.005, cor=-.31.

As we found a significant group effect on motor planning (CQscoreRT/Error) and partially significant correlations between planning and performance measures (sequence initiation time, movement time, and error rate), we conducted a mediation analysis to determine whether motor planning measures (CQscoreRT and CQscoreError) mediated the relationship between production variables (Sequence Initiation time, MT, Error rate) and a diagnosis of DCD.

### Error rate mediates the relationship between planning and DCD

We examined whether sequence planning as defined by CQscoreRT and CQscoreError mediates the relationship between sequence error rate and a DCD diagnosis. We found that group strongly predicted error (b = 29.11, p < .001), which was expected as there was a significant difference between groups in terms of sequence error rate. When controlling for both mediators (CQscoreRT, CQscoreError) we found that the direct effect of error on group was still strong, b = 23.30, p < .001. The total indirect effect when the mediators were combined was significant, b = .22, SE = .10, CI[.020, .42]. However, there was no effect of each specific mediator: CQscoreError, b = .089, SE = .075, CI[-.035, .26], or CQscoreRT, b = .13, SE = .079, CI[-.005, .30], although this neared significance. Therefore, this suggests that both planning measures significantly mediate the relationship between error rate and DCD.

We also investigated initiation time to see if it predicted DCD diagnosis when mediated by planning (CQscoreRT/Error). Initially, we found that group had a strong effect on initiation time, b = 78.95, p < .001. When controlling for both mediators, there was still a direct effect on group, b = 66.00, p < .001, suggesting that mediation is partial but not fully explained by planning. When examining each mediator individually, we found that neither CQscoreError, b = .056, SE = .058, CI[-.05, .19], nor CQscoreRT, b = .12, SE = .072, CI[-.012, .27] were significant individually, although CQscoreRT trended towards significance. Moreover, when both mediators were taken together, they did not significantly mediate the relationship between group and initiation time, b = .18, SE = .094, CI[-.0025, .37].

Finally, we investigated whether movement time predicted the relationship between planning and a DCD diagnosis when mediated by planning. As expected, we found that group has a strong effect on MT, b = 147.96, p < .001. When controlling for both mediators, we found that there was still a direct effect on group, b = 140.28, p = .001. However, we found that there was no significant effect of CQscoreRT, b = -.070. SE = .077, CI[-.26, .041], or CQscoreError, b = .11, SE = .086, CI[-.019, .31]. When the mediators were combined, there was still no significant indirect effect on MT, b = .042, SE = .094, CI[-.15, .23].

### DCD show working memory deficits but preserved inhibitory control

Next, we compared working memory scores between groups and examined whether working memory scores influenced movement planning/performance. Similar to other studies, we found that as task complexity increased, the DCD group had higher error rates compared to controls.

We observed an effect of group, F(1, 177) = 14.31, p < .001, no effect of nbacktype, F(2, 177) = .34, p = .71. An effect of age, F(1, 177) = 20.27, p < .001 but no interaction between group and nbacktype, F(2, 177) = .86, p = .42.

To assess inhibitory control, we used a go-nogo task with an 80% common and 20% rare condition and compared scores between groups (control and DCD). There was no effect of condition type, either 80% or 20%, F(1, 109) = .005, p =.94, and there was no interaction between group and condition type, F(1, 109) = .030, p = .86. This was also not affected by age, F(1,109) = .13, p = .72.

### Working memory score was not associated with motor planning

The correlations between CQscoreRT and working memory (nback3), rho=.015, p=.91, and CQscoreError and working memory (nback3), rho=.096, p=.46, reveal that working memory ability was unrelated to sequential planning (Figure 8).

We found that CQscoreRT and CQscoreError had a positive relationship, rho=.39, p=.002., showing that our measures of CQ correlated with each other. We found that after FDR correction, initiation time and CQscoreRT, rho=-.23, p=.14, and CQscoreError, rho=-.23, p=.14 were not significant. This shows that faster sequence initiation reflects a more pronounced CQ gradient.

Working memory score was not correlated with motor planning (CQscoreRT) for nback 1, rho=.20, p=.27, nback 2, rho=.096, p=.74, or for nback3, rho=.015, p=.97. CQscoreError was also not correlated with working memory score for nback 1, rho=.014, p=.97, nback2, rho=.087, p=.78 or for nback 3, rho=.096, p=.74, indicating that sequence planning is a separate mechanism to working memory.

Inhibition score for either the nogo80 and nogo20 was not associated with motor planning, for CQscoreRT and nogo80, rho=-.037, p=.97, CQscoreRT and nogo20, rho=-.11, p=.73. Or for CQscoreError and nogo80, rho=.014, p=.97, or CQscoreError and nogo20, rho=-.15, p=.49.

Furthermore, nback2, rho=-.39, p=.014, and nback3 were correlated with age, rho=-.39, p=.014. ADC score was correlated with movement time, rho=.34, p=.019, error rate, rho=.36, p=.005, nback3, rho=-.33, p=.046, and age, rho=.31, p=.036. This suggests that those who scored higher on the ADC checklist also had slower movement times, higher error rates and had poorer working memory, which supports previous research that working memory is impacted in DCD (Alloway, 2011).

## Discussion

Motor difficulties in individuals with DCD can affect everyday sequencing tasks such as tying shoelaces, handwriting, and typing. Although fluent and accurate coordination of sequential movements is essential for these activities, the mechanisms underlying such sequence coordination difficulties in DCD remain poorly understood.

Here, we demonstrate that the motor sequence planning difficulties observed in adults with DCD are fundamentally rooted in a reduced ability to pre-order upcoming movements. This deficit manifests as a compromised differentiation between successive sequence elements during preparation, leading to higher error rates during execution. Specifically, individuals with DCD exhibit smaller position-dependent differences, suggesting a greater neural overlap between planned elements. Furthermore, the DCD group were slower to initiate correctly sequences from memory and once initiated, produced the correct sequence at a slower pace. Previous work has shown that initiation time is correlated with the strength of the preordering (Mantziara et al., 2021), and our findings replicate this result. However, unlike error rate during sequence execution, the relationship between initiation time or movement time and DCD group was not significantly mediated by CQ score, suggesting that sequence timing deficits in DCD are not determined by the quality of the motor plan. Together, these findings imply that the pre-ordering of upcoming movements in a sequence underlies reduced sequence accuracy associated with DCD.

Our findings provide further support for the account of impaired motor planning in DCD and align with previous evidence demonstrating that motor planning deficits contribute to performance difficulties in this population (Adams et al., 2017; Bhoyroo et al., 2018, 2019; Fuelscher et al., 2016; Krajenbrink, Lust, Beckers, et al., 2021; Krajenbrink, Lust, & Steenbergen, 2021; Wilmut & Byrne, 2014). Studies using end-state comfort paradigms have shown that group differences become more pronounced as task complexity increases, with individuals with DCD exhibiting poorer planning performance. Our results extend this literature by demonstrating similar planning-related impairments in the context of a skilled sequential task.

Furthermore, we provide support for the theoretical account of motor impairment proposed by the internal modelling deficit (IMD) hypothesis, which suggests that individuals with DCD have difficulty generating and updating predictive internal models of action (Adams et al., 2014; Wilson, Smits-Engelsman, et al., 2017). Empirical support for the IMD hypothesis has largely come from studies employing motor imagery and mental rotation paradigms, such as hand rotation tasks (Barhoun et al., 2021; Hyde, 2014), which assesses the ability to mentally simulate movements. This body of work suggests that individuals with DCD have difficulty generating an accurate internal model, consistent with impaired motor planning (Adams et al., 2017; Barhoun et al., 2021; Hyde, 2014; Hyde et al., 2018; Kashuk et al., 2017). Our study builds on these findings by examining higher-level processes of motor planning. Specifically, we investigate feedforward planning indirectly through the structural organisation of sequential actions in skilled sequences, which requires the parallel competitive weighting of the upcoming items in a sequence, modelled as the parallel planning layer in computational CQ models (Burgess & Hitch, 2005; Houghton, 1990; Houghton & Hartley, 1996). Importantly, whereas previous work has inferred internal modelling deficits from performance on simulation-based tasks, our findings demonstrate that such deficits are evident in the real-time organisation of executable motor plans. By assessing movement pre-ordering, we show that in DCD, a more disorganised motor plan appears to reflect a weaker predictive model that is less effective for skilled motor performance.

To disentangle the mechanisms of working memory and sequence planning in DCD, we tested working memory capacity. Previously, a reduced working memory capacity has been found in children (Alloway, 2011), and adults with DCD (Wallinheimo & Gentle, 2024). Consistent with previous findings, we observed that adults with DCD had poorer working memory performance as task complexity increased (Wilson et al., 2013). Previous studies have found that working memory ability, particularly visuospatial memory, may play a role in the early stages of motor sequence learning (Hillman et al., 2024; Zuber et al., 2021). Therefore, we aimed to explore the relationship between working memory and motor sequence planning to see if this contributed to the impaired motor planning in DCD. We found that working memory was negatively correlated with ADC score, but working memory was not found to be correlated with sequence planning ability, as measured by CQ. This suggests that deficits in sequential planning are not driven by working memory capacity, but rather by a distinct mechanism underlying the motor coordination difficulties present in DCD. This dissociation could be because working memory and motor planning use distinct neural pathways (Eriksson et al., 2015; Wiestler et al., 2011).

In contrast, we observed no group-level differences in inhibitory control. This indicates that the observed motor planning deficits were entirely decoupled from general inhibitory constraints, suggesting that the graded finger-specific inhibition of upcoming movements during sequence preparation operates independently of global inhibitory control mechanisms (Duque et al., 2017; Labruna et al., 2014). Whilst some studies have reported inhibitory control deficits (He et al., 2018; Querne et al., 2008), a recent study using a stop signal task found no differences in stopping actions (Mayes et al., 2021). The authors concluded that this indicates a deficit in information processing rather than executive dysfunction, as the DCD group showed slower RTs (Mayes et al., 2021). Our findings reconcile these disparate accounts by demonstrating that while domain-general executive suppression may remain intact, the core deficit in DCD lies in the fine-grained computational processing required to pre-order and differentiate upcoming action representations during motor preparation.

Previous research has consistently linked motor planning deficits in DCD to dysfunction in brain regions such as the cerebellum (Zwicker et al., 2009), basal ganglia (Biotteau et al., 2016) and parietal regions (Kashiwagi et al., 2009; Zwicker et al., 2011). Our findings of impaired motor sequencing and planning in adults with DCD align with these studies. Notably, the cerebellum has been implicated in DCD, as it plays a crucial role in motor automatisation and sequencing (Brown-Lum & Zwicker, 2015; Biotteau et al., 2016). The cerebellum is thought to support fluent online prediction and the anticipatory motor control processes essential for effective motor planning (Kawato, 1999; Moberget & Ivry, 2016). In DCD, functional imaging studies have shown reduced activation in the parietal-cerebellar and fronto-parietal networks during motor tasks requiring planning and fine motor control. For example, during trail-tracing and joystick tracing tasks, both of which involve anticipating and executing precise movements, children with DCD show decreased activation in these networks (Kashiwagi et al., 2009; Zwicker et al., 2011). These findings suggest that impaired recruitment of these regions may underlie the reduced sequential planning in DCD. Additionally, the fronto-parietal network, particularly the left posterior parietal cortex (PPC) and postcentral gyrus, has been associated with sensorimotor integration and motor skill learning (Culham & Valyear, 2006; Debrabant et al., 2013; Querne et al., 2008; Wong et al., 2015). Lesions in the PPC, for instance, have been shown to result in visuomotor impairments, including deficits in movement sequencing and the generation of internal movement representations (Kashiwagi et al., 2009; Wong et al., 2015). Finally, our group has localised the pre-ordering of the upcoming sequence to the hippocampal and parahippocampal areas (Kornysheva et al., 2019; Yewbrey & Kornysheva, 2024), our current behavioural findings of a reduced pre-ordering of movements during planning raise the intriguing possibility that the hippocampus may be implicated in the sequencing deficits characteristic of DCD. Together, these studies support the theory that motor planning impairments in DCD are associated with functional disruptions in brain regions essential for coordinating, sequencing and predicting motor actions. Specifically, the hippocampal ordinal template may be weakened in adults with DCD, leading to a weaker sequence representation.

## Limitations

While this study offers novel insights into sequence planning in DCD, several limitations should be acknowledged. First, the DCD group was identified based on a self-reported diagnosis. Although the Adult Developmental Coordination Disorder Checklist (ADC) was also used to confirm key diagnostic criteria, such as symptom presence since childhood, it too relies on self-report, which may introduce subjectivity. To address Criterion D of the DSM-5, we collected information on educational attainment and screened for co-occurring neurological conditions to help ensure that the symptoms were not better explained by alternative diagnoses. Additionally, the study was conducted online, which, while allowing for broader participation, limits control over testing conditions. Although participants were given clear instructions and asked to complete the task in a quiet environment, we cannot verify full adherence to these guidelines. Finally, not all participants completed the working memory task: Control (N=42), DCD (N=20), and nogo task: Control (N=46), DCD (N=21).

Overall, our findings reveal that adults with DCD exhibit reduced pre-ordering of upcoming movements prior to execution, and this relationship between planning and DCD is mediated by error rate. This provides valuable insight into the specific motor planning mechanisms that underlie the performance differences observed, i.e. higher error rates, which were found to be distinct from working memory capacity or inhibitory control. Overall, these results indicate that the behavioural impairments defining adult DCD are fundamentally channelled through preparation constraints. Specifically, the capacity of elevated error rates to predict DCD group membership is mediated by a downstream failure in the fine-grained pre-ordering of upcoming movements.

## Author contributions

H.W., K.W. and K.K. designed research; H.W. performed research; H.W. and K.K. analysed the data; H.W. and K.K. wrote first paper draft; H.W., K.W. and K.K. edited the paper draft.

## Acknowledgements

This work was supported by Academy of Medical Sciences Springboard Award (SBF006/1052 to K.K.) and the UKRI Future Leaders Fellowship (MR/Y016467/1 to K.K).

